# Optimizing Plant Biofactories: Enhancing Recombinant Protein Production in *Nicotiana benthamiana* through Phytoplasma Effectors

**DOI:** 10.1101/2024.08.29.610350

**Authors:** Md Saifur Rahman, Marie-Claire Goulet, Dominique Michaud

**Affiliations:** Centre de recherche et d’innovation sur les végétaux, Faculté des Sciences de l’agriculture et de l’alimentation, Université Laval, Québec QC, Canada

**Keywords:** Molecular farming, *N. benthamiana*, phytoplasma effectors, recombinant protein, TENGU

## Abstract

Molecular farming, which utilizes plants as biofactories for recombinant protein production, offers an innovative and cost-effective alternative to traditional expression systems. Despite its advantages, plant-based production faces challenges such as low transgene expression and protein instability. Recent studies have highlighted the potential of *Nicotiana benthamiana* axillary stem leaves to enhance protein yield. This study explored the development of *N. benthamiana* lines expressing TENGU without signal peptide (T-SP), a phytoplasma effector known to induce plant dwarfism and increase shoot growth. TENGU and other effectors, such as SAP05 and SAP11, were introduced to create phenotypic variations that favor recombinant protein production. This study aimed to optimize these transgenic lines for increased biomass and protein yields by leveraging vertical farming conditions for scalable production. The results demonstrated significant improvements in leaf number, biomass, and five times more soluble protein content in T-SP lines compared to control lines, suggesting a promising approach for efficient molecular farming.

## 1. Introduction

The use of plants as biofactories for the production of recombinant proteins has emerged as a promising alternative to traditional expression systems in bacteria, yeast, and mammalian cells. This approach, known as molecular farming, leverages the natural capabilities of plants to produce complex proteins including vaccines and therapeutic antibodies in a cost-effective and scalable manner. Plant-based systems offer several advantages over traditional methods, including lower production costs, the ability to scale up quickly, and the potential to produce complex proteins with proper post-translational modifications. However, the efficiency of recombinant protein production in plants is often hindered by several challenges, including low transgene expression, protein instability, and suboptimal yield and quality of target proteins. Addressing these challenges is crucial to realizing the full potential of plant-based production systems.

Recent advances have highlighted the significant role of axillary stem leaves in enhancing recombinant protein yields in the transient protein expression host, *N. benthamiana*. Goulet et al. (2019) demonstrated that these leaves markedly contribute to protein yield, thereby presenting an opportunity to optimize expression systems for higher productivity (Goulet, et al., 2019). Building on this insight, our research focused on developing *N. benthamiana* lines that express TENGU, a phytoplasma polypeptide effector known to induce plant dwarfism and promote shoot growth in infected plants. TENGU, along with other effectors such as SAP04 and SAP11, holds potential in manipulating plant phenotypes to favor recombinant protein production (Carreón-Anguiano, Vila-Luna, Sáenz-Carbonell, & Canto-Canché, 2023).

Phytoplasmas are obligate parasitic bacteria that cause various plant diseases, leading to symptoms such as phyllody, virescence, and a bushy phenotype (Bertaccini, Oshima, Maejima, & Namba, 2019; Ermacora & Osler, 2019). These symptoms are primarily due to the action of virulence factors, such as TENGU, which interfere with plant hormone pathways, resulting in stunted growth and increased leaf proliferation. Minato et al. (2014) elucidated that TENGU downregulates the jasmonic acid and auxin pathways, leading to plant sterility and altered growth patterns (Minato, et al., 2014). Harnessing these unique properties of phytoplasma effectors can be strategically used to create dwarf plant biofactories with enhanced leaf biomass, potentially increasing the overall yield of recombinant proteins.

In this study, we aimed to develop stable transgenic lines of *N. benthamiana* that express TENGU and other phytoplasma effectors. Our short-term goals included inducing phenotypic variations that promote young leaf growth and increase heterologous protein yields in plants. In the long term, we sought to understand the physiological factors at the metabolic and proteomic levels that lead to an increased yield and quality of recombinant proteins. These insights will contribute to global efforts aimed at optimizing plant expression systems for large-scale production of efficient vaccines and therapeutic antibodies.

## 2. Materials and Methods

### 2.1 Experimental strategy

To achieve the goals of this study, we developed stable transgenic lines of *N. benthamiana* that express the phytoplasma effector TENGU. Phenotypic variations such as increased leaf growth and dwarfism were induced to enhance heterologous protein yields. A comprehensive analysis of physiological factors at the metabolic and proteomic levels was conducted to understand improvements in the yield and quality of recombinant proteins (Figure 1).

**Figure 1:**
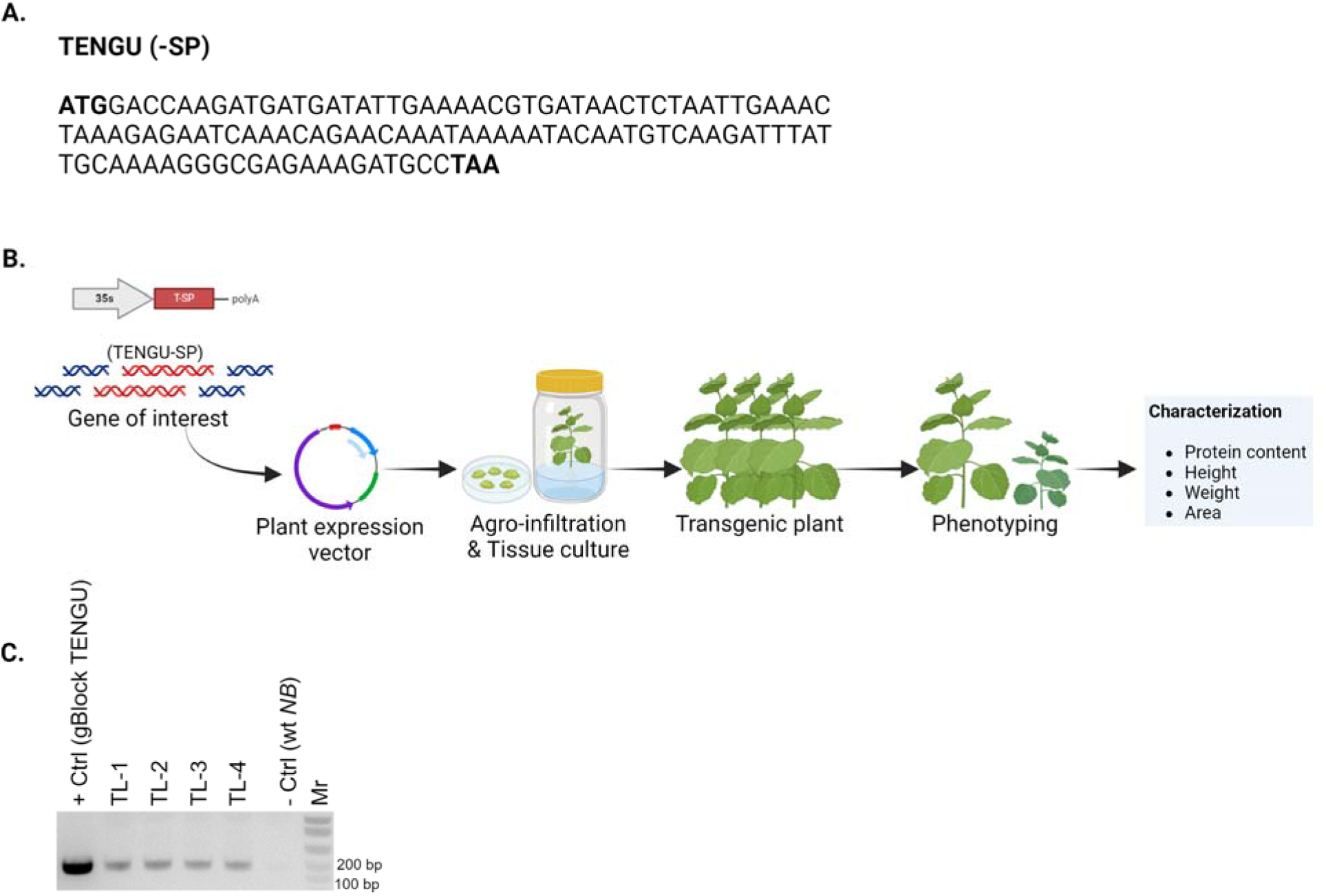
Stable transformation and PCR analysis of TENGU in *N. benthamiana*. (A) Coding sequence of the TENGU gene without signal peptide (B). Schematic representation of the stable transformation of TENGU in *N. benthamiana*. (C) Individual PCR for the confirmation of TENGU gene integration using gene specific primers in the selected positive transgenic plants. TL1, TL2, TL3, and TL4 independent *N. benthamiana* transgenic lines, + Ctrl : positive control (gBlock of TENGU), - Ctrl : Negative control (untransformed *Nb* plants), Mr: 1 kb plus marker (Invitrogen, USA).

### 2.2. Cloning, and plasmid constructs

Cloning procedures and plasmid constructs were performed according to methods described by Rahman et al. (2019) (Rahman, et al., 2021). Briefly, DNA sequences encoding the phytoplasma effectors TENGU, SAP05, and SAP11 were cloned into the binary vectors. These constructs were designed to facilitate stable integration and cloning using standard molecular Gateway cloning techniques. The constructs were confirmed by sequencing prior to plant transformation.

### 2.3. Transgenic plant development and tissue culture

Stable transformation of *N. benthamiana* was performed according to the protocol described by Stuttmann et al. (2019) (Ordon, Espenhahn, Kretschmer, & Stuttmann, 2019). *Agrobacterium tumefaciens* strain GV3101 harboring the transformation constructs was used to infect leaf explants. The explants were co-cultivated with Agrobacterium for two days on induction medium, followed by transfer to shoot induction medium (MS-II) containing kanamycin (100 mg/l) and cefotaxime (250 mg/l) for selection. Developing shoots were subsequently transferred to rooting medium (MS-III) to induce root formation. Once well developed, the transgenic shoots were acclimatized and transferred to the soil under high humidity conditions to establish mature plants.

The PCR analyses was performed using gene specific primers –

TNG-F: ATGGACCAAGATGATGATATTGAAAACG,

TNG-R: TTAGGCATCTTTCTCGCCCTTTTGC,

to confirm the presence of transgene in the putative homozygous transgenic plants both in T1 and T2 generation.

### 2.4. Plant material, growth conditions, and transgenic production

*N. benthamiana* plants used in this study were grown under controlled greenhouse conditions. Seeds were germinated in peat moss substrate and maintained at 24°C with a 16-hour light/8-hour dark photoperiod, ensuring a relative humidity of 60% and a light intensity of 130-150 µE m^-2^ s^-1^. The seedlings were transplanted into 350-ml pots and grown for an additional 3-5 weeks before transformation.

### 2.5. Phenotyping and characterization

Phenotyping and characterization of transgenic lines were performed according to the methods outlined by Goulet et al. (2019) (Goulet, et al., 2019). The assessed parameters included the number of leaves, plant height, leaf weight, and soluble protein content. Plant morphology was documented, and protein extraction was conducted using crude leaf extracts and standard biochemical assays to quantify the soluble protein content. Phenotypic traits were compared between T-SP lines and non-transformed control plants to determine the effect of TENGU expression on plant growth and protein yield.

### 2.6. Soluble protein extraction

Soluble protein content was determined from crude leaf extracts according to the methods described by Goulet et al. (2019) (Goulet, et al., 2019). Leaves were harvested and samples were taken from each plant. The samples were homogenized in extraction buffer (50 mM Tris-HCl, pH 7.5, 150 mM NaCl, 1 mM EDTA, and 0.1% Triton X-100) and centrifuged at 12,000 × g for 20 min at 4°C. The supernatant containing the soluble proteins was collected for analysis.

### 2.7. Protein quantification

The protein concentration in the leaf extracts was quantified using Bradford assay (Bradford, 1976). A standard curve was generated using bovine serum albumin (BSA), and the protein concentrations of the samples were determined by comparing their absorbance with the standard curve. The mean soluble protein content was calculated and reported as micrograms of protein per gram of fresh leaf weight.

### 2.8. Experimental design for productivity projection

#### Vertical farming setup

To evaluate the productivity of T-SP plants under vertical farming conditions, a growth chamber with multilevel shelving was set up to simulate a vertical farming environment. Each shelf in the chamber was equipped with LED lighting to provide a uniform light intensity across all levels. The light intensity was maintained at 150 µE m^-2^ s^-1^ with a 16-hour light/8-hour dark photoperiod. Temperature and humidity were controlled to match the conditions described for greenhouse growth.

#### Plant density and spacing

Plants were arranged to optimize the use of vertical space. Control and T-SP plants were grown in separate sections of the chamber. The control section contained approximately 8 plants per cubic meter (m^3^), while the T-SP section was optimized to contain up to 45 plants per m^3^ due to their compact growth form.

### 2.9. Statistical analysis

All experiments were conducted with a minimum of six biological replicates. Data were statistically analyzed using analysis of variance (ANOVA), followed by Tukey’s multiple comparison test to determine significant differences between means. *P-value* of 0.05 was used as the threshold for statistical significance.

## 3. Results and Discussion

### 3.1. Partial characterization of T-SP *N. benthamiana* lines

Partial characterization of T-SP *N. benthamiana* lines revealed significant differences in plant morphology and protein content compared to the control lines. As depicted in Figure 2A, T-SP plants exhibited a higher number of leaves than the control plants. Specifically, the T-SP lines had an average of 45 leaves, whereas the control lines had approximately eight leaves. This substantial increase in leaf number is a direct consequence of TENGU expression, which induces a bushy phenotype and promotes shoot growth (Sugawara, et al., 2013). In terms of plant height (Figure 2B), the T-SP lines showed a significantly reduced stature, with an average height of 13.6 ± 1.1 cm compared to the control plants’ height of 22.3 ± 3.7 cm. This dwarf phenotype is characteristic of TENGU’s effect on plant growth, which downregulates hormone pathways such as jasmonic acid and auxin, resulting in shorter but more proliferative plants (Bertaccini, et al., 2019). Leaf weight was also markedly increased in the T-SP lines (Figure 2C). The average leaf weight of T-SP plants was 418.0 g, significantly higher than the 65.8 g observed in control plants. This increase in biomass is beneficial for recombinant protein production as it directly correlates with the potential yield of proteins harvested from plants. The most crucial aspect for molecular farming, the mean soluble protein content in crude leaf extracts, was significantly higher in the T-SP lines (Figure 2D). The T-SP lines exhibited a notable increase in soluble protein content compared with the control plants. This increase is likely due to the enhanced biomass and number of leaves, which provide more material for protein extraction. The observed increase in protein content underscores the potential of T-SP plants to serve as effective biofactories for recombinant protein production (Bertaccini, et al., 2019)

**Figure 2:**
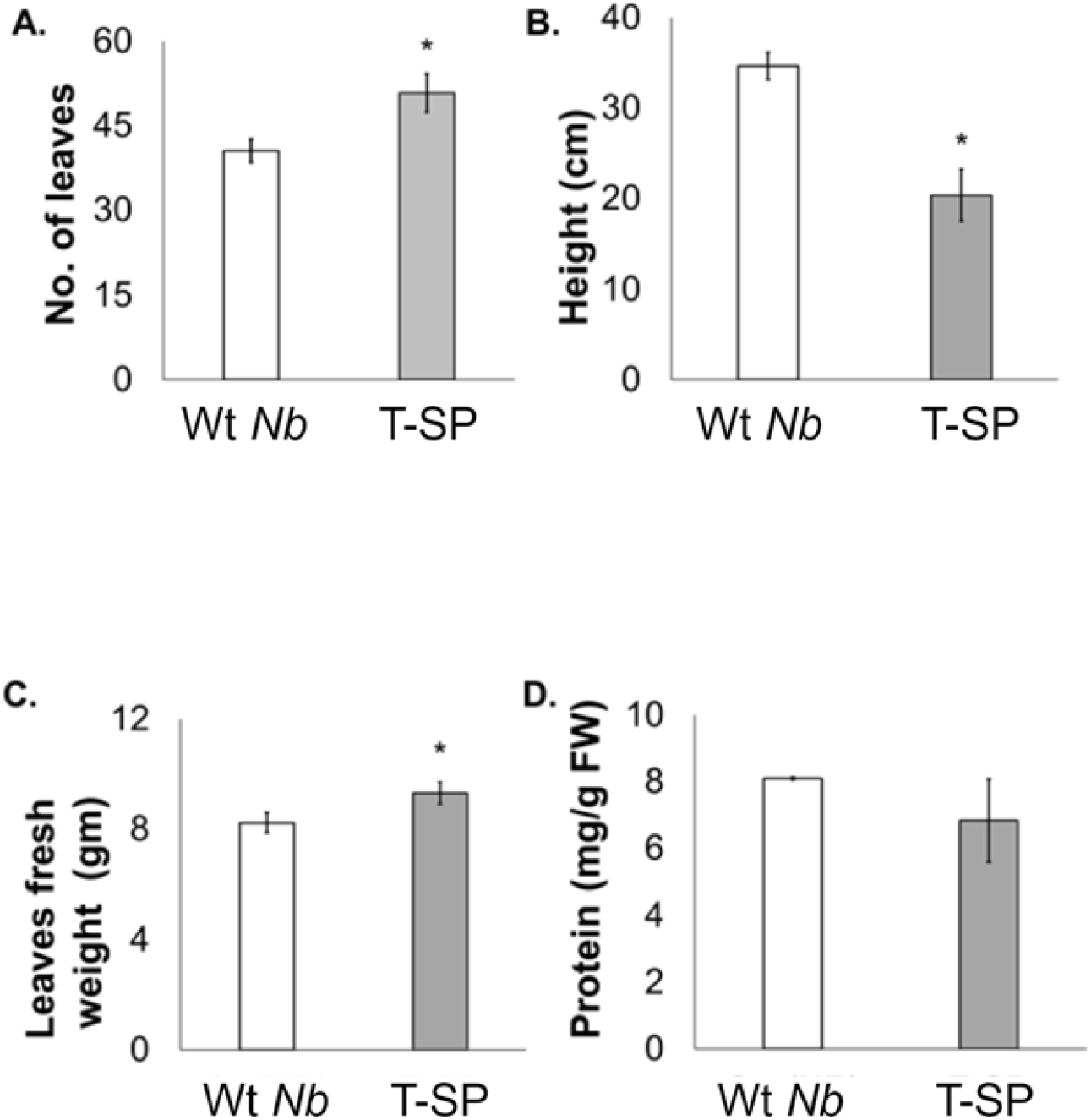
Partial characterization of T-SP *N. benthamiana* line. (A) Number of leaves. (B) Plant height. (C) Leaf weight is expressed as the mean of four biological replicates ± SD. (D) Mean soluble protein content in crude leaf extracts from six biological replicates ± SD. * Significance at *p-value* < 0.05.

### 3.2. Projected productivity under vertical farming conditions

Figure 3 presents a projection of the productivity of 6-week-old control and TENGU-SP plants per cubic meter (m^3^) under multilevel vertical farming conditions. This projection assumes a similar yield of recombinant protein per milligram (mg) of leaf tissue across the different lines. The projection illustrates a stark contrast in the number of cultivable plants per cubic meter between the control and the TENGU-SP lines. Control plants, owing to their taller stature, can accommodate approximately 8 plants per m^3^, whereas the compact growth habit of TENGU-SP plants allows for a much denser cultivation, supporting up to 45 plants per m^3^. When comparing the fresh leaf weight per m^3^, the T-SP lines again demonstrated a significant advantage (Figure 3). The T-SP plants yielded a total fresh leaf weight of 418.0 g per m^3^, compared to the control plants (65.8 g. This considerable increase in leaf biomass translates directly to an enhanced potential for recombinant protein production, assuming that the protein yield per milligram of leaf tissue remains consistent. The projected productivity data further suggested that TENGU-SP plants could yield several times more protein than control plants within the same cultivation volume. This enhanced productivity is pivotal for the economic viability of molecular farming, as it maximizes the use of available space and resources, thereby reducing the overall production costs (Bertaccini, et al., 2019).

**Figure 3:**
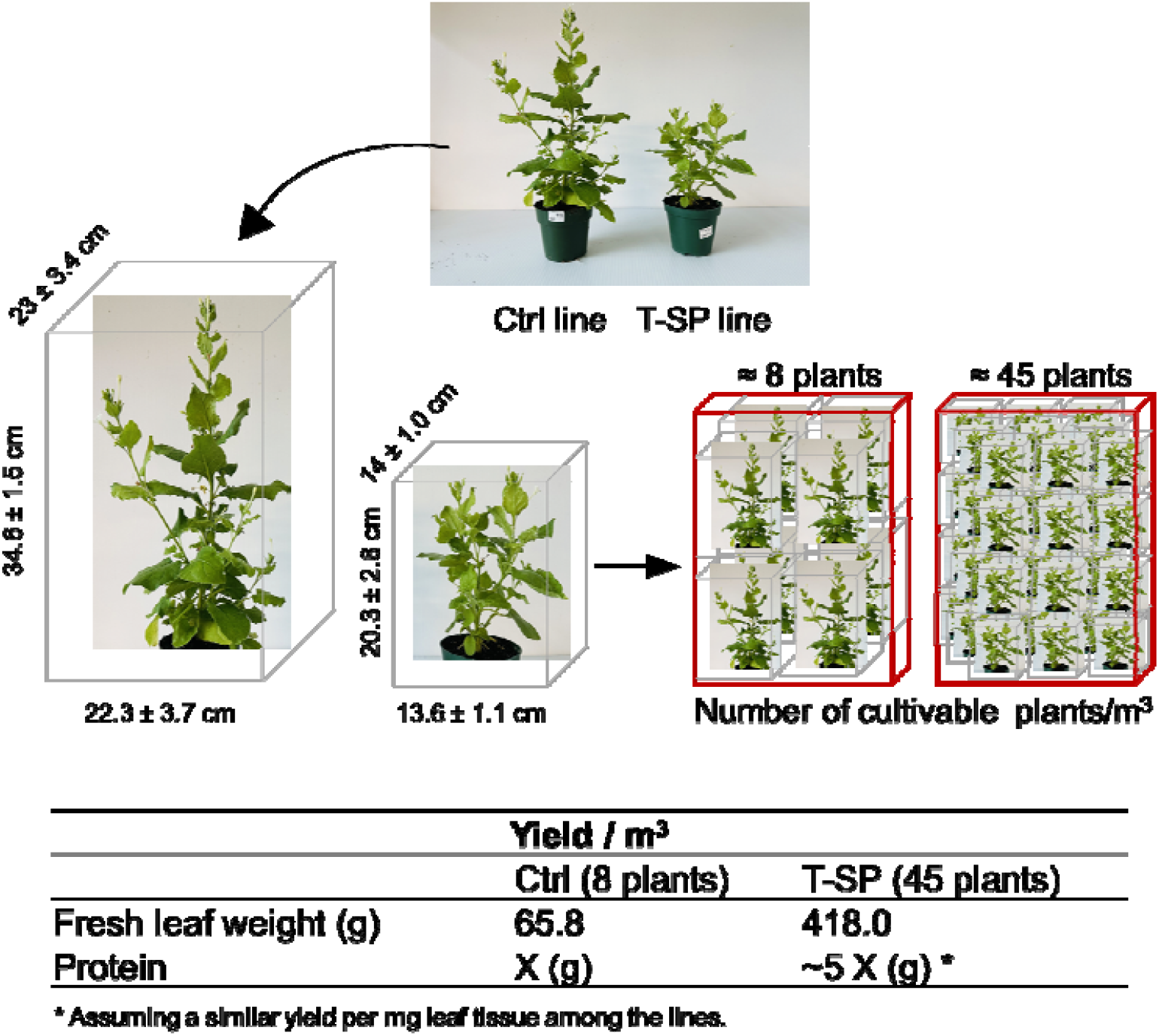
Projected productivity under vertical farming conditions. Projected productivity of 6-week-old control and TENGU-SP plants per m^3^ under multi-level vertical farming conditions, assuming a similar yield of recombinant protein per mg leaf tissue.

The results collectively highlight the significant potential of using T-SP *N. benthamiana* lines for molecular farming applications. The increased leaf number, reduced plant height, and enhanced biomass observed in the T-SP plants contributed to a more efficient production system for recombinant proteins. The dwarf phenotype induced by TENGU expression aligns with the goal of optimizing plant architecture for dense cultivation setups such as vertical farming. The ability to cultivate more plants per unit volume not only increases the overall biomass but also enhances the potential yield of recombinant proteins, making the production process more cost-effective and scalable (Bertaccini, et al., 2019; Sugawara, et al., 2013). The significant increase in soluble protein content observed in the T-SP plants further underscores the practical benefits of using these transgenic lines. A higher protein content in the leaf tissue means that more recombinant protein can be extracted from the same amount of plant material, thereby improving the efficiency of the production process (Bertaccini, et al., 2019; Sugawara, et al., 2013). These findings support the hypothesis that manipulating plant phenotypes through the expression of phytoplasma effectors, such as TENGU, can lead to substantial improvements in recombinant protein production. Further enhancements in plant biofactory systems can be achieved by understanding and leveraging the molecular mechanisms underlying these phenotypic changes (Arya, Rookes, Cahill, & Lenka, 2020; Buyel, Stöger, & Bortesi, 2021; Singh, 2023).

To further advance the findings of this project on optimizing plant biofactories through the expression of phytoplasma effectors like TENGU in *N. benthamiana*, several key experiments are necessary:

#### Full characterization of T-SP lines

A comprehensive analysis involving molecular, physiological, and morphological characterizations of T-SP plants is required. This would include detailed assessments of growth patterns, reproductive capabilities, and metabolic profiling to better understand the modifications induced by TENGU expression.

#### Selection and confirmation of homozygous lines

It is imperative to develop and select homozygous lines for the introduced traits to ensure consistency and stability of the phenotypes across generations. This process will also help in assessing any potential positional effects of the transgene insertion, which could influence expression levels and phenotypic outcomes.

#### Experimental therapeutic antigen production

Utilizing the optimized transgenic lines, production trials for specific recombinant proteins, such as vaccines or therapeutic antigens, should be conducted. This will validate the practical applications of these biofactories in producing high-value proteins under controlled conditions.

#### Scale-up under vertical farming conditions

To evaluate the scalability of this approach, experiments should be conducted in a vertical farming setup. This will assess the biomass and recombinant protein yield per unit area, providing insights into commercial viability and operational logistics.

## Conclusion

The development of T-SP *N. benthamiana* lines represents a promising approach for overcoming the challenges of low transgene expression and protein instability in plant-based production systems. The significant improvements in biomass and protein yield observed in this study highlight the potential of these transgenic lines to revolutionize molecular farming, paving the way for the more efficient and cost-effective production of vaccines and therapeutic antibodies.

## Ethical Compliance

N/A

## Data Access Statement

The data presented in this study are available on request from the corresponding author.

## Author Contributions

SR, MG, and DM contributed to the design and implementation of the research, SR to the analysis of the results and to the writing of the manuscript. SR and DM conceived the original and supervised the project.

## References

Arya, S. S., Rookes, J. E., Cahill, D. M., & Lenka, S. K. (2020). Next-generation metabolic engineering approaches towards development of plant cell suspension cultures as specialized metabolite producing biofactories. Biotechnology Advances, 45, 107635.

Bertaccini, A., Oshima, K., Maejima, K., & Namba, S. (2019). Phytoplasma effectors and pathogenicity factors. Phytoplasmas: Plant Pathogenic Bacteria-III: Genomics, Host Pathogen Interactions and Diagnosis, 17–34.

Bradford, M. M. (1976). A rapid and sensitive method for the quantitation of microgram quantities of protein utilizing the principle of protein-dye binding. Analytical biochemistry, 72, 248–254.

Buyel, J. F., Stöger, E., & Bortesi, L. (2021). Targeted genome editing of plants and plant cells for biomanufacturing. Transgenic Research, 30, 401–426.

Carreón-Anguiano, K. G., Vila-Luna, S. E., Sáenz-Carbonell, L., & Canto-Canché, B. (2023). Novel Insights into Phytoplasma Effectors. Horticulturae, 9, 1228.

Ermacora, P., & Osler, R. (2019). Symptoms of phytoplasma diseases. Phytoplasmas: methods and protocols, 53–67.

Goulet, M.-C., Gaudreau, L., Gagné, M., Maltais, A.-M., Laliberté, A.-C., Éthier, G., Bechtold, N., Martel, M., D’Aoust, M.-A., & Gosselin, A. (2019). Production of biopharmaceuticals in Nicotiana benthamiana—axillary stem growth as a key determinant of total protein yield. Frontiers in Plant Science, 10, 735.

Minato, N., Himeno, M., Hoshi, A., Maejima, K., Komatsu, K., Takebayashi, Y., Kasahara, H., Yusa, A., Yamaji, Y., & Oshima, K. (2014). The phytoplasmal virulence factor TENGU causes plant sterility by downregulating of the jasmonic acid and auxin pathways. Scientific reports, 4, 7399.

Ordon, J., Espenhahn, H., Kretschmer, C., & Stuttmann, J. (2019). Stable transformation of Nicotiana benthamiana.

Rahman, M. S., Madina, M. H., Plourde, M. B., Dos Santos, K. C. G., Huang, X., Zhang, Y., Laliberté, J.-F., & Germain, H. (2021). The fungal effector Mlp37347 alters plasmodesmata fluxes and enhances susceptibility to pathogen. Microorganisms, 9, 1232.

Singh, H. (2023). Enhancement of Plant Secondary Metabolites by Genetic Manipulation. In Genetic Manipulation of Secondary Metabolites in Medicinal Plant (pp. 59–90): Springer.

Sugawara, K., Honma, Y., Komatsu, K., Himeno, M., Oshima, K., & Namba, S. (2013). The alteration of plant morphology by small peptides released from the proteolytic processing of the bacterial peptide TENGU. Plant Physiology, 162, 2005–2014.

